# De novo design of alpha-beta repeat proteins

**DOI:** 10.1101/2024.06.15.590358

**Authors:** Dmitri Zorine, David Baker

## Abstract

Proteins composed of a single structural unit tandemly repeated multiple times carry out a wide range of functions in biology. There has hence been considerable interest in designing such repeat proteins; previous approaches have employed strict constraints on secondary structure types and relative geometries, and most characterized designs either mimic a known natural topology, adhere closely to a parametric helical bundle architecture, or exploit very short repetitive sequences. Here, we describe Rosetta-based and deep learning hallucination methods for generating novel repeat protein architectures featuring mixed alpha-helix and beta-strand topologies, and 25 new highly stable alpha-beta proteins designed using these methods. We find that incorporation of terminal caps which prevent beta strand mediated intermolecular interactions increases the solubility and monomericity of individual designs as well as overall design success rate.

## Introduction

Repeat proteins can serve to connect, join, arrange, and bind diverse biomolecular targets, both in nature and in bioengineering applications. These functions emerge from a combination of globally repetitive architecture and diversity in local sequence. A wide variety of design and engineering approaches have been employed to add functionality to ^1–3^ and extend the range of geometries sampled^4^ by natural repeat proteins. While modifying and mimicking existing natural protein backbones has yielded a number of notable successes in design ^4–8^, these approaches are fundamentally limited by the robustness of the native protein to mutation, and are limited in design space to solutions close to the parent protein. On the other extreme, a broad geometric diversity of repeat proteins has been generated with all-helical geometry ^9,10^, but there has been less progress with extended loops or beta-strand containing structures. A particular challenge with beta sheet containing structures is the potential for aggregation through exposed beta strands at the termini. Natural repeat proteins containing beta strands often have irregular helical segments as capping elements; it is unclear whether such elements are also necessary for design.

Here we describe the development and characterization of general methods for designing beta-sheet-containing repeat proteins. We explored both Rosetta ^11^ and deep learning based methods for generating such structures, and the role of explicitly designed capping elements to prevent aggregation.

## Results

### Rosetta based design approach

We generated Alpha-Beta Repeat (ABR) proteins using Rosetta Remodel fragment assembly ^12,13^ and Rosetta sequence design (Figure 1A). We performed remodel trajectories based on “blueprint files” with fixed secondary structure fragment lengths within a repeat unit. We tandemly repeated the exact backbone structure for each repeat unit, with varied distance constraints between specified repeat positions. Initial blueprints were chosen to broadly mimic similar natural proteins, such as LRR and TLR repeats with between two and four helical heptads and strand lengths between four and nine residues. Loop lengths were sampled initially at very permissive ranges of two to nine residues, and later constrained to two or three residues, after low sampling efficiency was observed. Blueprint files were designed with a specific order of secondary structure fragment type in the following order: strand, loop, helix, loop. In order to improve beta strand alignment *in silico*, fragment sizes were based on “structural rules” for the design of beta-containing proteins ^14^ for loops under 5 residues.

**Figure 1.**
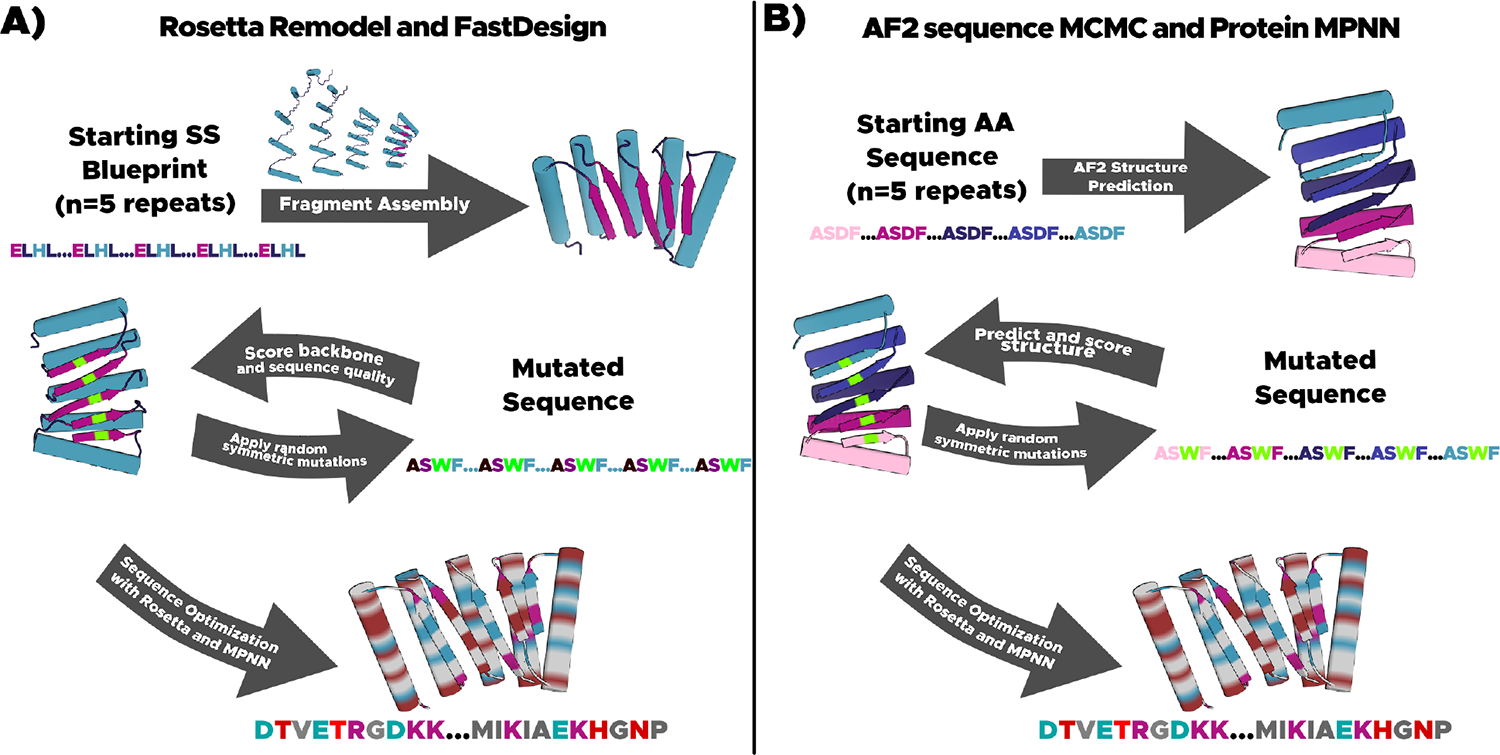
Computational methods for designing Alpha-Beta Repeat Protein **A)** Repeat protein design using the Rosetta Protein Design Suite. Fragment assembly is conducted with a blueprint specifying secondary structure sizes, lengths, and order. Backbones are screened for plausible geometry, and core positions are designed with symmetric FastDesign. Surfaces are then designed with asymmetric FastDesign, to decrease surface hydrophobic residues frequency. B) Repeat protein design using MCMC sequence random walk with AlphaFold2 prediction. Backbone models are generated by prediction from a tandem repeated sequence. Predicted structures are evaluated for prediction confidence (plDDT, pTM) and a mixture of alpha-helices and beta-strands (DSSP). Hydrophobic core residues are designed with Protein MPNN, and surface positions are designed with Rosetta FastDesign.

To enforce the beta-strand pairing between repeat units, we introduced atomic distance constraints between secondary structure fragments as well as backbone carbon atoms of strand regions during the fragment assembly trajectory. We ran trajectories using a variety of different constraint regimes: weak constraints between alpha-carbon (Ca) atoms at strand repeat positions, constraints between Ca and beta-carbon (Cb), and constraints between Ca, Cb, as well as helix-helix distances. Ideal Ca and Cb distances were sampled between 4.5 and 5.5 Angstroms while inter helix distances were sampled between 8 and 12 angstroms. We screened for designability of the fragment assembled backbone models based on a combination of fragment quality and backbone geometry metrics ^15^ and presence of beta-strands by secondary structure prediction (using the Dictionary of Secondary Structure of Proteins method,DSSP, within Rosetta^16^). These trajectories produced backbone models with a twisted parallel beta-sheet.

### Deep Learning based design approach

For DL design of protein backbones we conducted tandem repeat hallucination Markov Chain Monte Carlo (MCMC) sequence searches guided by AlphaFold2 (AF2) predictions as previously described ^17,18^, but with constraints on predicted secondary structure content rather than superhelical parameters or global architecture (Figure 1B), and without filtering for closed ring shaped structures. We carried out sequence space Markov Chain Monte Carlo (MCMC) optimization with sequence length L and number of repeating units N individually specified for each trajectory (Figure 2). Trajectories were seeded with a sequence predicted to fold (at any confidence) to a structure with alpha-helices and beta sheets, then tandemly repeated N times. These initial sequences were generated in one of three ways: 1) from the pool of Rosetta-based designs, 2) by using a repeating Leucine-Leucine-Alanine sequence for helices, poly-Valine for sheets, and repeating glycine-serine for loops, or 3) by collecting the repeat sequences generated from initial trajectories using the first two methods. Monte Carlo search was guided by a weighted linear combination of AF2 model confidence (predicted LDDT), predicted TMScore (pTM) and sequence proportion of alpha helices and beta sheets (ranging between 0.3 and 0.6 for each secondary structure respectively); sequence substitutions were accepted or rejected based on the standard Metropolis criterion.

**Fig 2.**
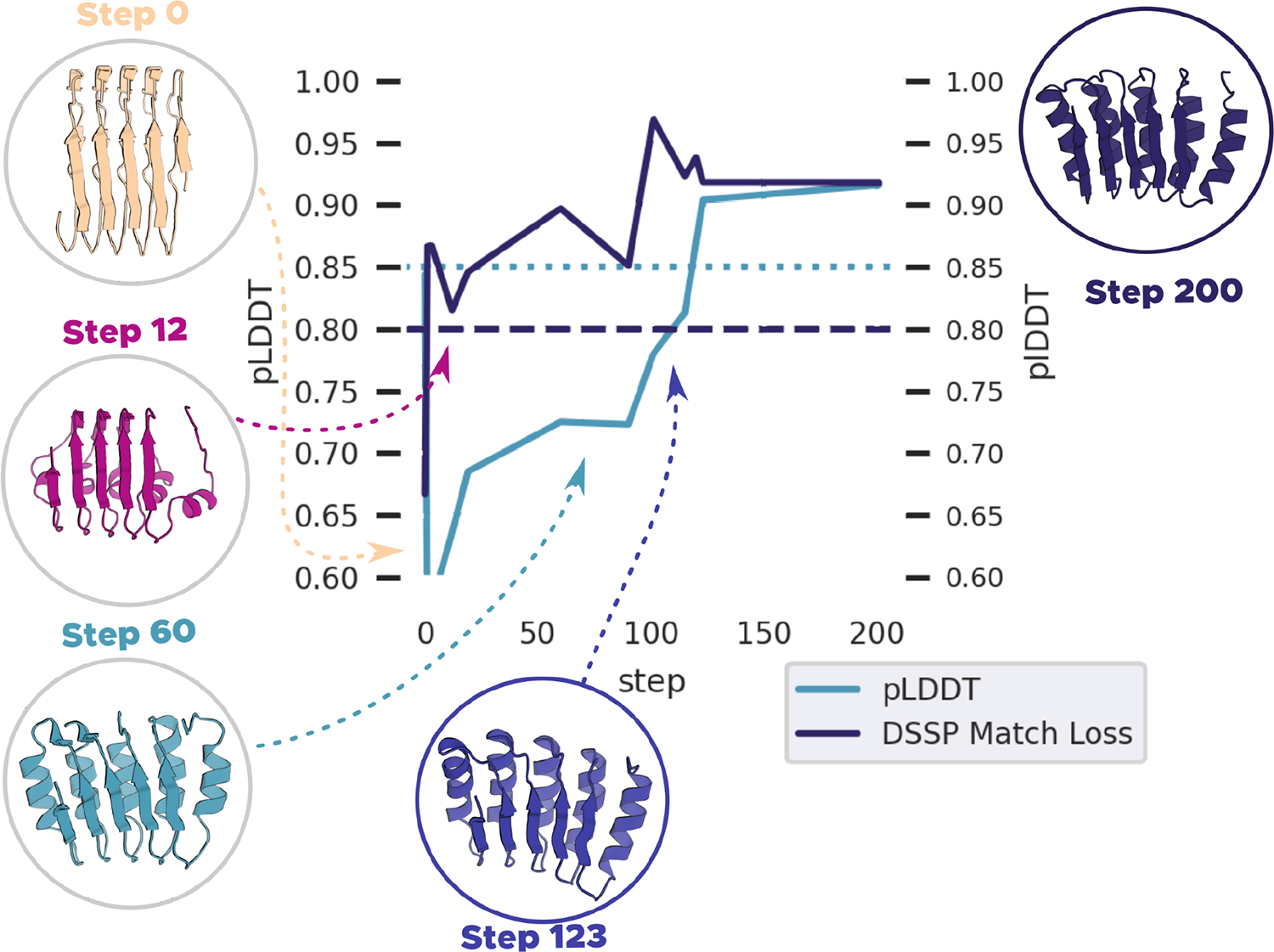
Example of Hallucination Trajectory A sample hallucination trajectory of MonteCarlo accepted models shown with corresponding traces of two losses (plDDT and DSSP match loss). “DSSP match” measures the agreement of the actual secondary structure (Helix, Sheet, Loop) and target secondary structure, and plDDT reports on prediction confidence. Dashed and dotted horizontal lines indicate minimal “designable” cutoffs for further design steps.

### Capping Feature Design

We reasoned we could improve our success rates through designed helical capping features. Native beta-strand-containing proteins almost always have such caps, and rapidly folding helix-containing segments could potentially block slower forming intermolecular strand-strand interactions. We selected as a test system a design with a reversible, concentration dependent soluble aggregation mode that had much higher beta character signal on CD at high concentrations (Figure S5). We hypothesized that if the transition from a folded mixed fold to a lower energy solenoid or amyloid aggregate was driving aggregation of our designs, low contact order helical capping features might provide a kinetic barrier to prevent this transition.

Helical capping features were designed with Rosetta remodel, and sequence design was conducted with distance constraints anchoring the alpha-helical capping features in place at the termini of the repetitive region. Constraints were enforced between the cap terminus and the nearest residue in the proximal repeat unit after each cycle of Rosetta FastRelax. The quality of capping was determined by changes in the Surface Aggregation Propensity (SAP) ^19^ score measured with and without the cap in the final models. Of the models with lowest SAP scores, the model with the largest difference between SAP score measured with and without the cap was chosen for characterization. AF2 prediction of caps sequence designed with Rosetta deviated considerably from the original design model, with capping regions often not contacting or protecting the terminal regions; we found this method unreliable for capping feature design. We thus chose final design sequences through 3 iterations of MPNN, AF2 prediction, and Rosetta FastRelax to obtain. The quality of capping was determined by changes in the Surface Aggregation Propensity (SAP) ^19^ surface score measured with and without the cap in the final models. Of the models with lowest SAP scores, the model with the largest difference between SAP score measured with and without the cap was chosen for characterization. We were unable to generate sequences for which the capping features remained at the design site for the Rosetta design group of models using standard Rosetta distance constraints and Layer Design, and therefore solely characterized capping features on DL designed repeat proteins.

### Sequence Design

We conducted sequence design with either RosettaFast Design or ProteinMPNN. For Rosetta FastDesign, we applied strict sequence symmetry for core residue positions and unconstrained sequence design for solvent accessible regions. We selected designs based on standard Rosetta metrics (see methods for details). For MPNN design, no sequence symmetry was applied. We conducted MPNN design by iterating over MPNN sequence design, Rosetta FastRelax, and AF2 prediction. We selected MPNN designed sequences based on AF2 prediction by filtering on pLDDT and pTM, and further narrowed the pool by selecting on Rosetta metrics (see methods for details).

### Experimental characterization

Synthetic genes encoding 85 selected Rosetta designs and 47 selected DL designs with helical endcaps were obtained, and the proteins expressed and purified from E coli. 57 designs were expressed solubly, and 35 displayed monomeric peaks on SEC. However, many were purified at low yield, displayed concentration dependent aggregation, or precipitated out of solution over the course of a few days at refrigerated temperatures. These results are summarized in table 1. While the various Rosetta distance constraint design schemes did reduce the sampling time required to produce computational successes (and therefore increased the number of designs we were able to characterize), they came at a trade off of a lower number of viable repeat protein sequences purified under laboratory conditions.

**Table 1.** Successful Design Hits by Backbone Generation Protocol. The fraction of models characterized by soluble, monodisperse expression as assayed by distinct peak with no shoulder on SEC, at a retention volume outside the column void volume. DL design produced a dramatically higher success rate among investigated models than constrained fragment assembly. Percentages shown are rounded to the nearest whole integer.

| Design Protocol | Number of Models Tested | Count Designs Soluble and Monodisperse | Percent Designs Soluble and Monodisperse |
| --- | --- | --- | --- |
| **All Methodologies** | **132** | **35** | **27%** |
| Weak Constraints | 45 | 7 | 16% |
| CA & CB CST | 21 | 3 | 15% |
| CA,CB & Helix CST | 19 | 0 | 0 |
| DL Design | 47 | 25 | 43% |

The DL designs had a much higher success rate than those produced with conventional Rosetta Design procedures when considered alone. Over half of them expressed solubly, and many more had monodisperse peaks on SEC at a useful yield for circular dichroism characterization and crystallographic screens (Table 1). They expressed at high yields, even in small scale 50 ml cultures, they remained soluble overnight after IMAC but before SEC, and showed refolding character on CD, even after heating to 95C.

Given this difference between groups, we sought to clarify the relative impact of Protein MPNN sequence design and the explicit design of helical capping features in the DL designed molecules. In order to separately investigate these two effects, we chose 6 diverse designs from the DL design stream which we classified as “highly soluble” (they yielded soluble material eluting at a retention volume consistent with a monomeric or dimeric species on SEC). In 5 of the 6 investigated cases, deletion of both caps resulted in less soluble yield than a capped design (Figure 4). In the case where a fully uncapped protein performed best, the second best performer was a singly capped model with nearly identical yield.

In order to account for exposed hydrophobics resulting from a direct sequence deletion, we also applied a whole-structure MPNN compensatory redesign to each of our 18 cap deletion designs. Once again, 5 out of 6 designs performed best with caps. Full results are summarized in Figure 4. Solubility and expression was quantified through paired expression and purification of test models. The peak absorbance at 230 nm during SEC was taken as a metric of usable yield. In each case, the yield was normalized to the highest peak observed, so the best model was taken as a normalized absorbance of 1.0, and each other model was quantified as a fraction of this amount. Peaks corresponding to soluble aggregate (retention ml of under 12 ml for the S200 columns used) were not considered as usable yield.

### Fusion to Designed Heterodimers

Following the successful design, expression and characterization of the base repeat proteins, we explored their use as scaffolds by fusing them along the edge strands to beta strand containing proteins. We chose the monomeric subunits of reversibly forming designed heterodimers (LHDs) to enable incorporation of our designs into higher order assemblies and to add functional domains through fusion to the complementary LHD subunit. LHD subunits have been fused to DHRs ^21^, but this process is limited by location of helices on the LHD subunit, and results in fusions which twist dramatically away from the plane along which the molecules dimerize (Supplementary Fig S4), We docked our ABR repeat proteins with LHDs along terminal strands with a simple strand template alignment script and designed backbone connections with Protein Inpainting ^20^. We then designed these models as described above with MPNN, AF2, and FastRelax: conserving the original heterodimer interface, but allowing mutations to accommodate the new fusion. The resulting fusions expressed solubly in E coli, and the purified proteins co-eluted with the complementary LHD subunit on SEC (Supplementary Fig S4).

## Discussion

Using a combination of Rosetta blueprint design and deep learning based hallucination, we were able to robustly design new repeat proteins with beta strand containing repeats. While previous attempts to design repeat proteins with extensive strand pairing were restricted to highly constrained topologies derived from hydrogen bonding patterns found in nature, our methodology was able to discover a wide variety of repeat proteins using a relatively unconstrained search strategy. We found that terminal helices on designed, extended beta structures often improved soluble yields, with a single helix at one terminus sometimes improving expression or purification by an order of magnitude. We expect our capping procedure can be generally applied to improve solubility of designed proteins with terminal beta strands. Our designed ABRs and their ability to accommodate rigid fusion with LHD adapters should allow construction of a wide variety of beta sheet containing assemblies in a manner analogous to the way that helical repeat proteins have contributed to a wide diversity of alpha helical assemblies ^22,23^.

## Methods

### Computational Design Methods

#### AF2 Hallucination Trajectories

We conducted MCMC sequence searches scored by quality and confidence of AlphaFold2 (AF2) predictions. Optimization starts from a specification, within each trajectory, of the length L and number N of repeating units. A portion of trajectories were seeded with a sequence predicted to fold (at any confidence) to a structure with alpha-helices and beta sheets, then tandemly repeated N times. The remainder were seeded with random sequences. Only a small fraction of the random sequences converged to a folded structure containing both alpha-helices and beta-sheets, and these sequences were then used to seed future trajectories. Overall, thousands of ABR backbones with plDDT of 0.9 or greater were produced from several thousand computational trajectories (successful trajectories produce many plausible backbone models). We used a scaling mutation rate starting from 5 per step and decreasing down to 1 (based on how many steps had elapsed).

#### DL Assisted Sequence Design

We conducted sequence design by combining ProteinMPNN with Rosetta FastRelax for 5 cycles. Each cycle starts with ProteinMPNN used to predict a sequence, then a model of the protein was obtained via AF2. This model was relaxed with the default Rosetta FastRelax. The cycle was then repeated an additional four times. Models were chosen by the criteria of an RMSD of 2 or lower to the initial AF2 hallucination prediction, and plDDT of 0.8 or better. We then redesigned surface positions with Rosetta LayerDesign, selecting only solvent accessible positions for design. Final sequences were then predicted again by AF2, and models chosen for characterization again by a threshold of RMSD of 2 or lower to the initial AF2 hallucination prediction, and plDDT of 0.8 or better.

#### Cap Design

We designed minimal helical caps through the combination of Rosetta Remodel and ProteinMPNN sequence prediction.. ABR sequences with caps designed through Rosetta tools alone consistently failed to form capping features, hence, we chose to investigate the impact of these capping features with the DL designed pool. Cap backbones were designed with Remodel, specifying loop lengths of two or three residues, and helix lengths sampled from 10 to 20 residues. Capping was evaluated by a decrease in SAP score of the residues of the terminal repeat region when considered with versus without the cap structure. A sequence for the whole structure was then predicted with the ProteinMPNN and FastRelax procedure described above. Structures with at least one capping feature predicted contacting the terminal repeat hydrophobic residues were considered “capped”.

#### Protein Expression and Purification

We prepared recombinant protein for both Rosetta designs and DL based designs in the same fashion. We obtained cloned gene plasmids and Golden Gate compatible gene sequences encoding the designs with a cleavable tag (SNAC tag ^24^) with downstream hexahistidine tag. In the case of Golden Gate cloning, final sequences were confirmed either by intact protein mass spectrometry via an electrospray ionization time of flight (ESI-TOF) of expressed protein or directly by colony sequencing of a single colony used for expression by Azenta. All plasmids used contained a sequence conferring Kanamycin resistance. Plasmids were transformed into BL-21 DE3 E. coli and expressed protein via auto-induction. Autoinduction was conducted in 50 ml cultures with 6 hour outgrowth at 37℃ from either a 50% glycerol stock prepared from an overnight culture or from an agar plate colony, followed by a 24 hour expression at 18℃ before culture harvest.

TerrificBroth (TB) with 5052 and Kanamycin Sulfate (as a selection factor) was used as auto-induction media. Similar yields were observed between different expressions from glycerol stocks, single colony inoculations, and small variations in expression time at 18℃. Cell cultures were harvested from media by centrifugation in 50ml conical tubes, and cell pellets were frozen at −20C until purification.

Lysis was conducted by sonication on ice in 20 mM Tris, 100mM NaCl, and 50 mM Imidazole buffer at pH 8 with protease cocktail tablets used according to manufacturer instructions. Sonicated lysate was chilled before centrifugation. After centrifugation at 20,000g for 40 minutes, lysate supernatant was purified by immobilized metal affinity chromatography (IMAC) with nickel NTA resin (brand) by batch binding at 4℃ followed by elution with 20mM Tris, 100 mM NaCl and 500 mM imidazole. Eluates were stored at room temperature overnight before fast protein liquid chromatography (FPLC). On the subsequent day, eluates were centrifuged at minimum 20,000g for 12 minutes, and supernatant was [assed through a 0.2 micron filter before FPLC. Size Exclusion Chromatography (SEC) was conducted with S200 and S75 columns from (brand etc) based on expected molecular weight and resolution appropriate for a particular preparation.

Circular Dichroism (CD) data were obtained with a Jasco J-1500. We conducted a thermal temperature melt with temperature ranges of 25℃ baseline temperature, a mid temperature of either 55℃ or 65℃ and a high temperature of 95℃. An “annealed” measurement was also taken after the sample was cooled down to 25℃. Traces were found to be consistent with a structure having both alpha-helical and beta-strand character (figure 3).

**Figure 3.**
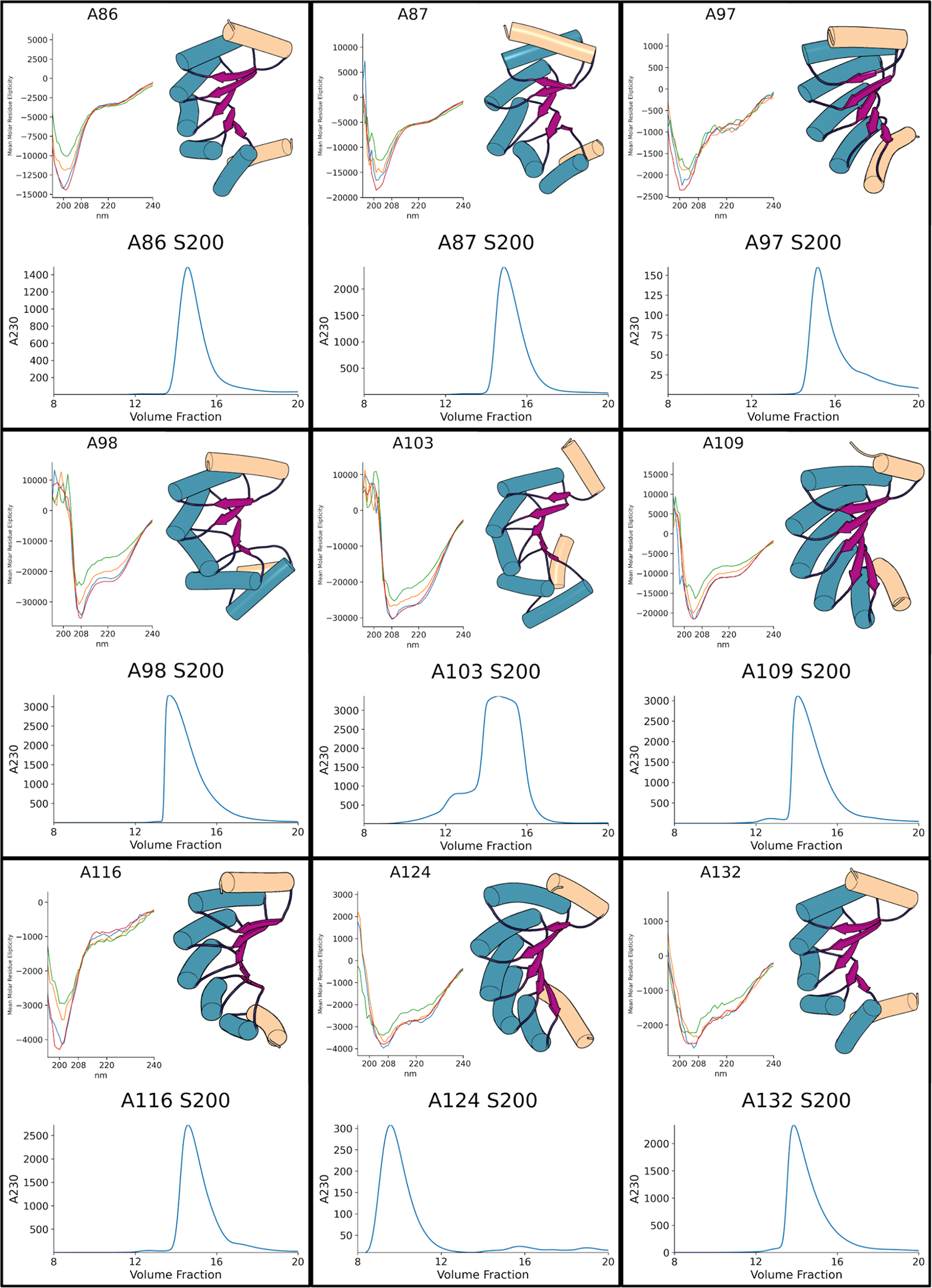
Experimental characterization. CD spectra were measured at 25C and 95C followed by a final spectrum recorded at 25C to assess refolding. A middle temperature trace is shown for each at either 65C or 55C. The blue and red traces correspond to 25C at the beginning of the melt and after refolding, respectively. The orange trace shows the scan at either 65C or 55C, the green trace shows 95C. Capping features are colored in tan. Repeat unit helices are colored blue, loops dark purple, and strands in violet.

**Figure 4.**
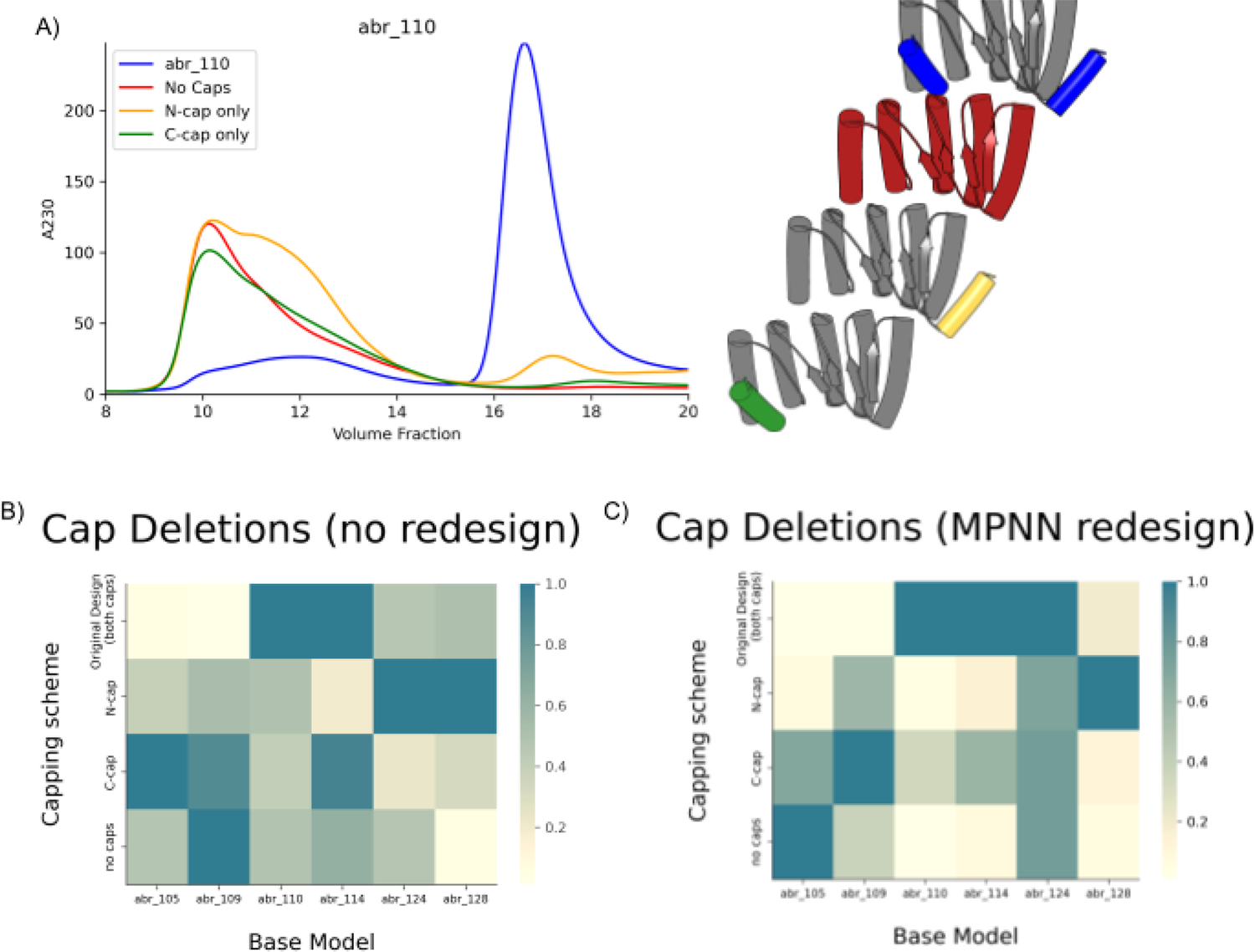
Contribution of capping elements to soluble expresssion A) Different cap configurations for 6 designs were expressed and purified in parallel. Soluble yield is estimated by the highest point in the SEC trace, in the expected retention volume range for the construct as a monomer. Left panel, SEC traces; Right panel, models of designs with and without caps using the same color scheme. B) & C) Heatmaps of soluble yield for different cap deletions. Yield is normalized to the highest yielding construct in the design pool. Models in B are sequence deletion mutants, while models in C) are redesigned with MPNN after deletion. In both cases, all but one backbone model performed best with at least one cap.

## Disclaimers and Copyright

Figures S3, 5, and 6 are reprinted by permission with or without modification from D. Zorine’s PhD thesis ^25^

## Acknowledgements

The authors would like to acknowledge Linna An, Derrick R. Hicks, Lukas Milles, Alexis Courbet, Basile Wicky, Florian Praetorius, and Danny Sahtoe for help in adapting tools, methods, and design strategies from related projects for this work. The authors would like to acknowledge the use of unpublished python packages during design iteration: including “npose”, “homog”, and “getpy” created by Brian Coventry, Will Sheffler, and Adam Moyer respectively. These packages are imported in updated hallucination code.

The authors would also like to acknowledge the work of the IPD’s core research facilities in providing support and reagents for this paper. Mila Lamb and Xinting Li ran and obtained the intact protein mass spectrometry data to validate the identity of these proteins. Stacey Gerben, Analisa Murray, and Piper Heine produced SNAC-cleaved protein for use in experiments and crystallography trials.

This work was supported by a grant from the Department of Defense (DOD-0001039633), a grant from the National Institute on Aging (5U19AG065156.), a gift from the Washington Research Foundation, and the Audacious Project at the Institute for Protein Design.

This research used resources of the National Energy Research Scientific Computing Center, which is supported by the Office of Science of the U.S. Department of Energy under Contract No. DE-AC02-05CH11231.

## Supplementary info 1 ABR Rosetta Design

### FastDesign and Sequence design procedures

Rosetta sequence design process involved a number of stages with modifications to the default FastDesign parameters. The FastDesign procedure can be broken down into three consecutive design protocols, included as code in [**Supplementary Code XX**]. Each protocol comprises MCMC sequence sampling and backbone angle minimization. During this process mutation permutations are rapidly sampled through “packing” steps, where coarse grained atom coordinate trees assess relative self-compatibility of various sequences, and minimization is conducted via explicit gradient descent of Rosetta Energy via changes to backbone torsion angles and residue chi angles. Detailed explanation of the three packing/minimization protocols follow. Default FastDesign is described in^26^.

### “Sandpig” QuickPack

A swift packing and minimization round was conducted with only the set of residues with one-letter codes SANDPVG. This procedure is meant to relax poly-leucine and poly-alanine backbones into geometries more compatible with FastDesign sampling, while keeping backbone hydrogen bonds satisfied. The rationale for this step was to improve compatibility between backbones derived from fragment assembly under the centroid score function and sequences with full-atom scoring. Following “sandpig” packing, glycine and proline positions introduced at this step remained fixed within the structure for the remainder of the design process. This protocol is inspired by graduate work previous published from our group ^27^.

### Symmetric Layer Design

The symmetric sequence design protocol comprises the “LayerDesign’’ framework in Rosetta. In summary, residues were categorized into core, surface, and boundary positions based on their number of “neighbors”: residues whose “neighbor atoms” (typically the beta-carbon) are within a mutual threshold distance. High neighbor count residues are considered “core”, low count are considered “surface”, and values between the two are designated as “boundary”. In the LayerDesign protocol, certain residue identities are forbidden for use in packing. Core positions forbid use of polar residues, while surface positions forbid the use of hydrophobic residues (except proline). In addition, our layer design scheme featured discrimination between secondary structure features, and forbade use of “helix breaking” residues in helices (Asparagine, Aspartate, proline) and bulky residues in loop regions.

Additionally, a custom relaxscript file with adjusted reference weights was used to reduce heavy bias within Rosetta to assign histidine to every position in parallel beta strands. Standard reference weights were used during final FastRelax and scoring.

Symmetric layer design was conducted for 4 cycles of packing and minimization. The “approx_buried_unsat” term was used during packing to penalize buried, unsaturated polar residue head groups and backbone polar heavy atoms.

This LayerDesign protocol was conducted with the repeat symmetric constraints in Rosetta, where residue identity and geometry between repeat units is exactly fixed and accounted for in scoring.

### Surface Design

Surface positions were designed following a procedure similar to that of symmetric layer design, with the key difference being the absence of symmetric constraints. Only surface residues and boundary residues on strands were marked as designable. This allowed for the modification of sequences at terminal repeats to break symmetry and enhance residue diversity on the strand surfaces. Symmetric sequence constraints in Rosetta FastDesign typically resulted in sheets composed entirely of a single residue identity, observed as histidine and serine during preliminary testing.

### Scoring and Filtering

A variety of score terms in rosetta were applied to discriminate promising models for investigation.Rosetta designs were chosen by selecting the models with the lowest number of buried, unsaturated heavy atoms and the best scores among “worst9mer” (a fragment quality metric), sidechain shape complementarity, and rosetta score per residue. For each of the 7 characterization rounds, a range 50,000-100,000 candidate backbone models were generated, of which only approximately 500-2,500 were suitable for design. Models failing initial screens were deleted to conserve disk space and not investigated further. Between 5 and 25 sequence design trajectories were run on each backbone, as computational resources allowed. Rather than firm score thresholds, the best models were selected from the pool based on these metrics. Model sequences for investigation were chosen from the top quintile of each metric.

## Supplementary info 2 AF2 MCMC and Protein MPNN Design

### Backbone Generation Protocol

A process of sequence hallucination was used to explore mixed alpha- and beta-character backbones through Markov Chain Monte Carlo (MCMC) sampling of sequence/structure pairs. This hallucination script is included in supplementary code https://github.com/dmitropher/af2_multistate_hallucination and is only slightly modified from the procedure used in other related works ^17,18^.

Initial models were constructed by AF2 prediction of a selected tandem repeated sequence with a length (L) ranging from 18 to 45. This sequence was either chosen at random, taken from a repeat unit of an existing Rosetta design sequence (as designed in Supplementary Info 1), or seeded manually. Repeat unit sequences from successful trajectories were also used for follow-up trajectories to increase sampling. Manual sequence seeds were chosen to emulate the “blueprint” strategy from Rosetta design: helical stretches, loops, and strands were defined with residues that have high statistical propensity for those sequences in nature. Helical segments were seeded with the repeat sequence “AAL”, loops with “GGS” and strands with “VS”.

At each iteration, a quality score was derived from random mutations that were assessed by their effects on the backbone predicted from that sequence by AF2. Mutations were applied in tandem to each residue with the same position within the repeat unit (1,1+L,1+2L etc were all mutated from some type X to some other type Z). This structure underwent evaluation based on an evenly weighted linear combination of presenting both alpha-helices and beta-sheets, as well as favorable pLDDT and pTMscore. A “Fuzzy Fraction of Secondary Structure” score was derived by seeding the trajectory with a desired fraction of each type of secondary structure (helix, loop, and strand) in the final prediction, and a score from 0 to 1 (with 0 being the worst, 1 being the best) was given for adherence to that fraction of secondary structure in the final model. The score was an evenly weighted linear combination of logistically rescaled difference in fraction from the given values. Small deviations (10% or less) were considered to be a deviation of 0% in the final evaluation (the “Fuzz” factor). The objective of the protocol was to generate structures with both alpha-helices and beta-sheets, rather than some exact proportion of these structures. Mutations were introduced at a rate of 1-7 per MCMC step, with more mutations at the trajectory’s outset and fewer as it progressed.

### Design of Helical Caps

Helical caps were designed at the N and C termini using Rosetta remodel. Helices of lengths between 14 and 25 residues were sampled and prepended or appended to structures along with a connecting loop of 2 or 3 residues. Backbones containing these helical caps were sequence designed asymmetrically using MPNN.

### Sequence Design

Following the generation of initial ABR scaffolds, and addition of helical caps, ProteinMPNN was employed for designing sequences as asymmetric monomeric proteins. The Rosetta sequence design suite was employed for a redesign of the ABR surface residues, using the “surface design” protocol as described in Supplementary Info 1. Final sequences were then predicted with AF2 to ensure fidelity, with final designs being selected for having less than 2.5 RMSD to the initial prediction and plddt of 0.8 and TMscore of 0.65 or greater as evaluated by AF2.

## Supplementary Data 1 - ABR Additional SEC and CD Data

**Figure S.1.**
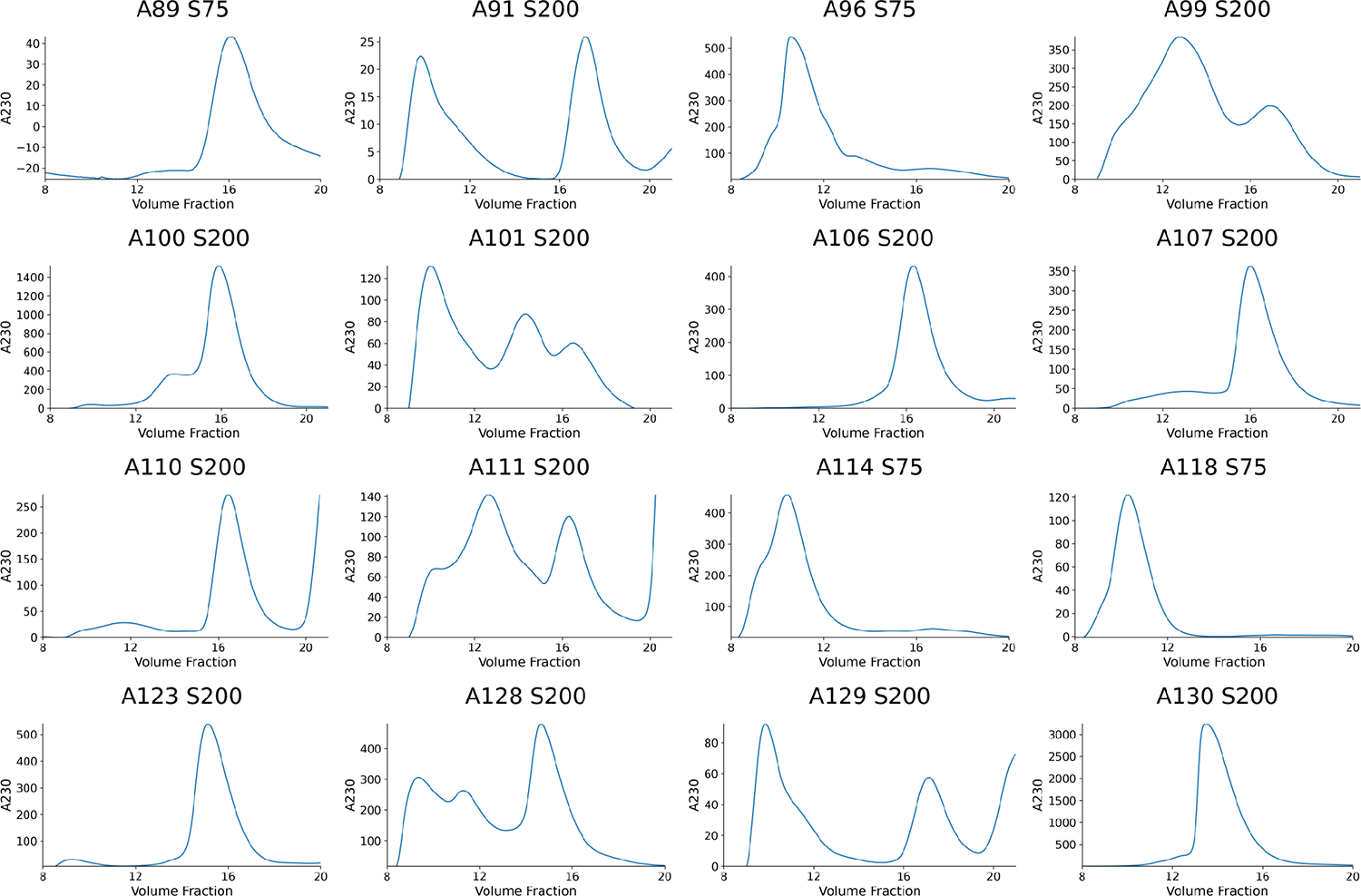
Selected Additional ABR SEC traces Select purification and analytical traces from the column indicated for each sample. Each protein was selected either because the peak appears at the expected retention volume, or a high-concentration peak is present at a retention volume greater than the column void volume. These designs were considered as having a monodisperse fraction for the purposes of Table 1.

**Figure S. 2.**
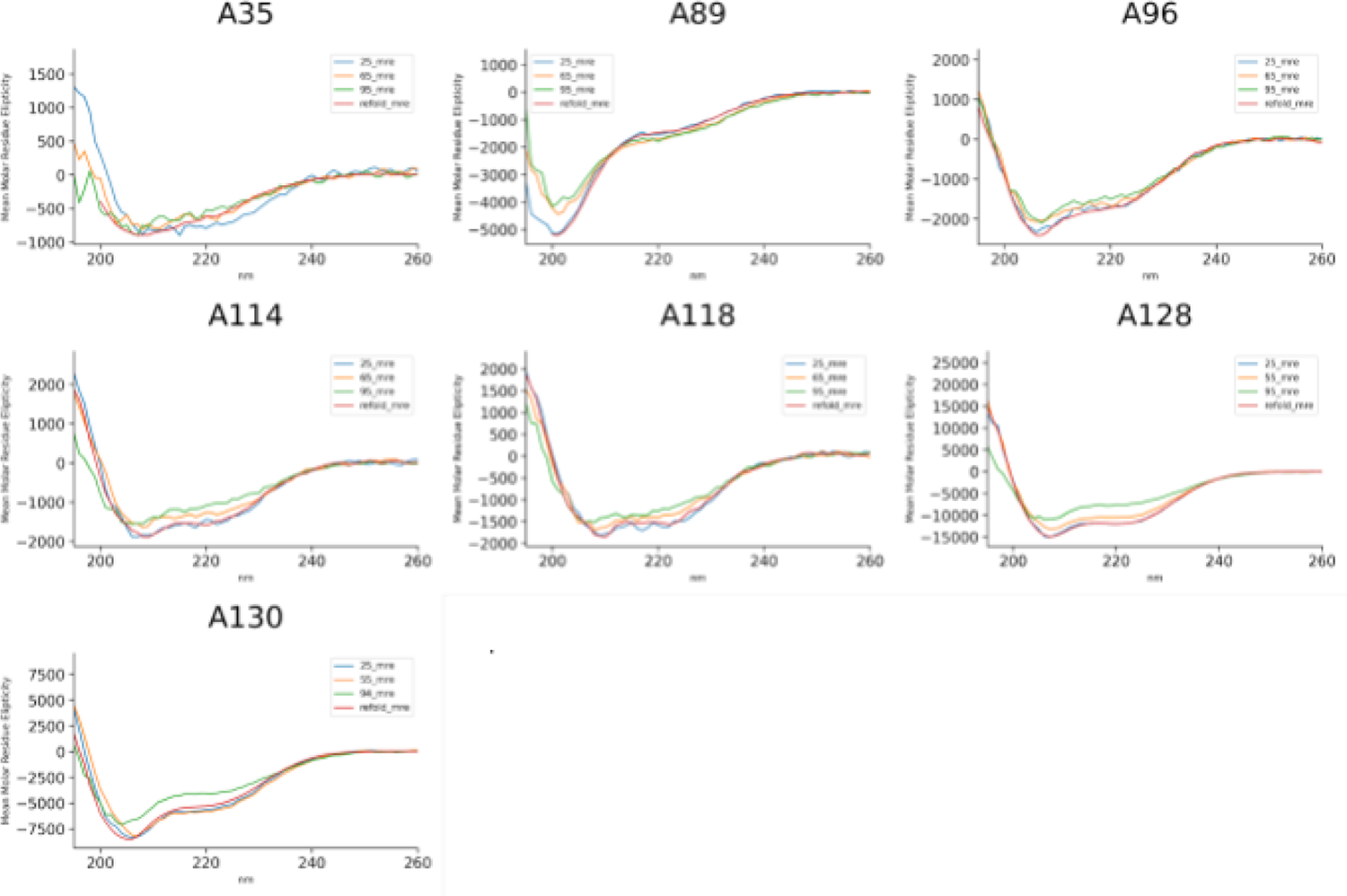
Additional CD melt traces Additional CD scans conducted during a thermal melt from 25C to 95C. Traces are presented at 25C (blue), 55C or 65C as noted (orange), and 95C (green), as well as a scan taken after cooling to 25C (red) post-melt

**Figure S.3.**
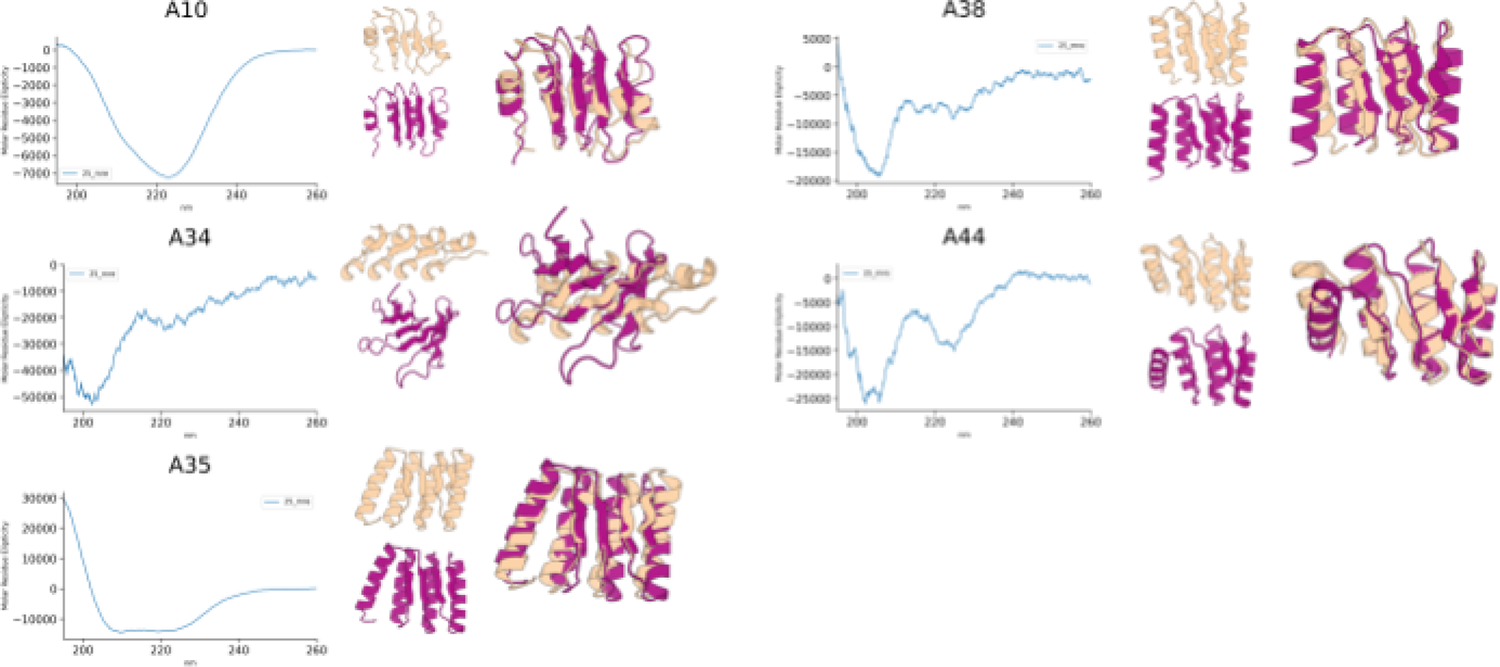
CD Scans Rosetta Designs CD Scans for selected Rosetta designs. At right, the design model is shown in tan while an AF2 prediction is shown in violet. An overlay is shown of both models. Low yields and lower solubility in the low-salt buffer resulted in poor data collection for A34,38, and 44, with features lacking sufficient definition for secondary structure characterization

**Figure S.4.**
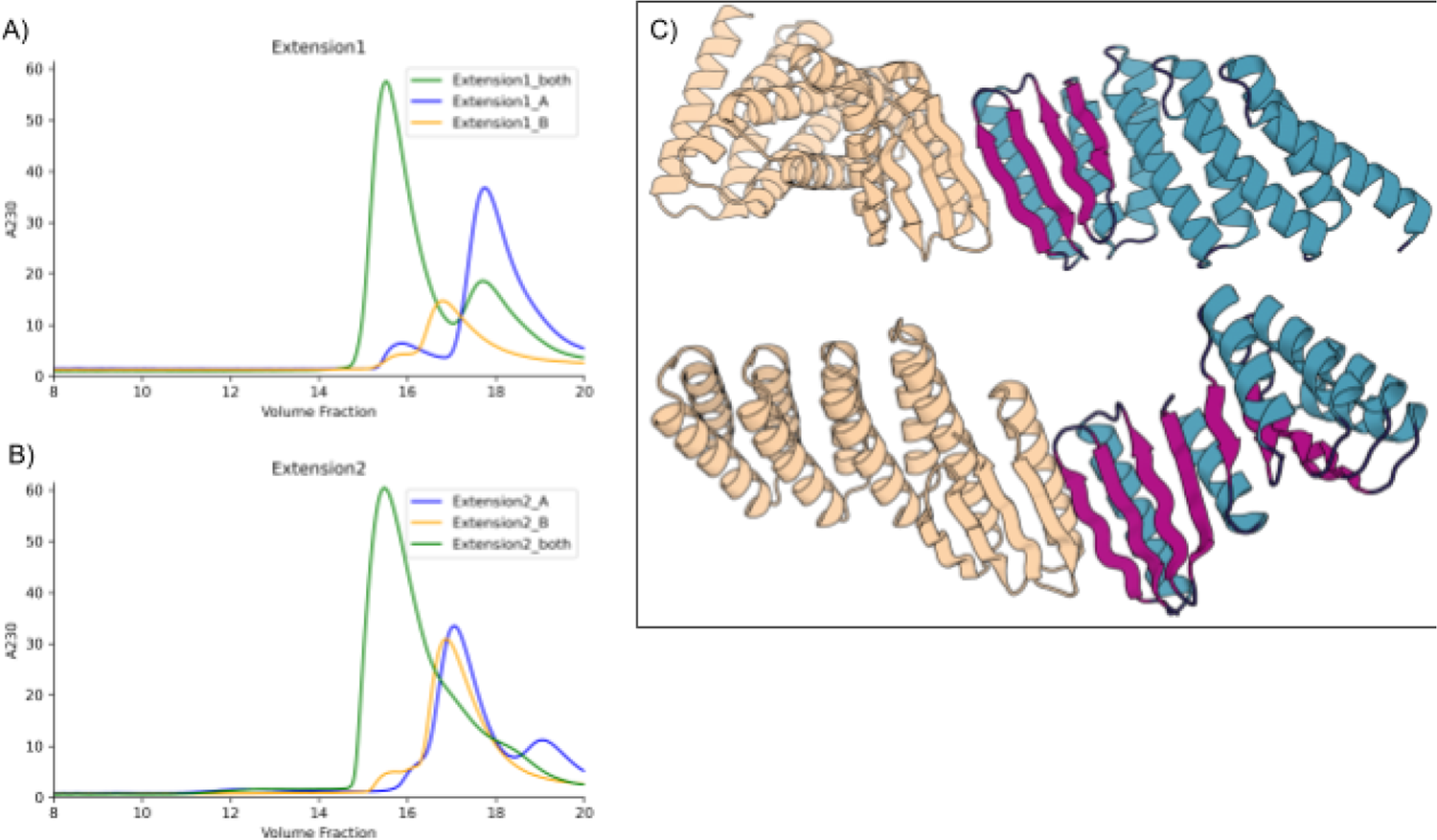
Panels A) and B) show SEC co-migration of abr extended dimers. Panel C contrasts a previously published, DHR extended, LHD with our new extension scheme. Both schemes allow for fusion to a functional heterodimer. Our new fusions extend the sheet region and fuse at different relative orientations than are available to DHR fusions.

**Figure S.5.**
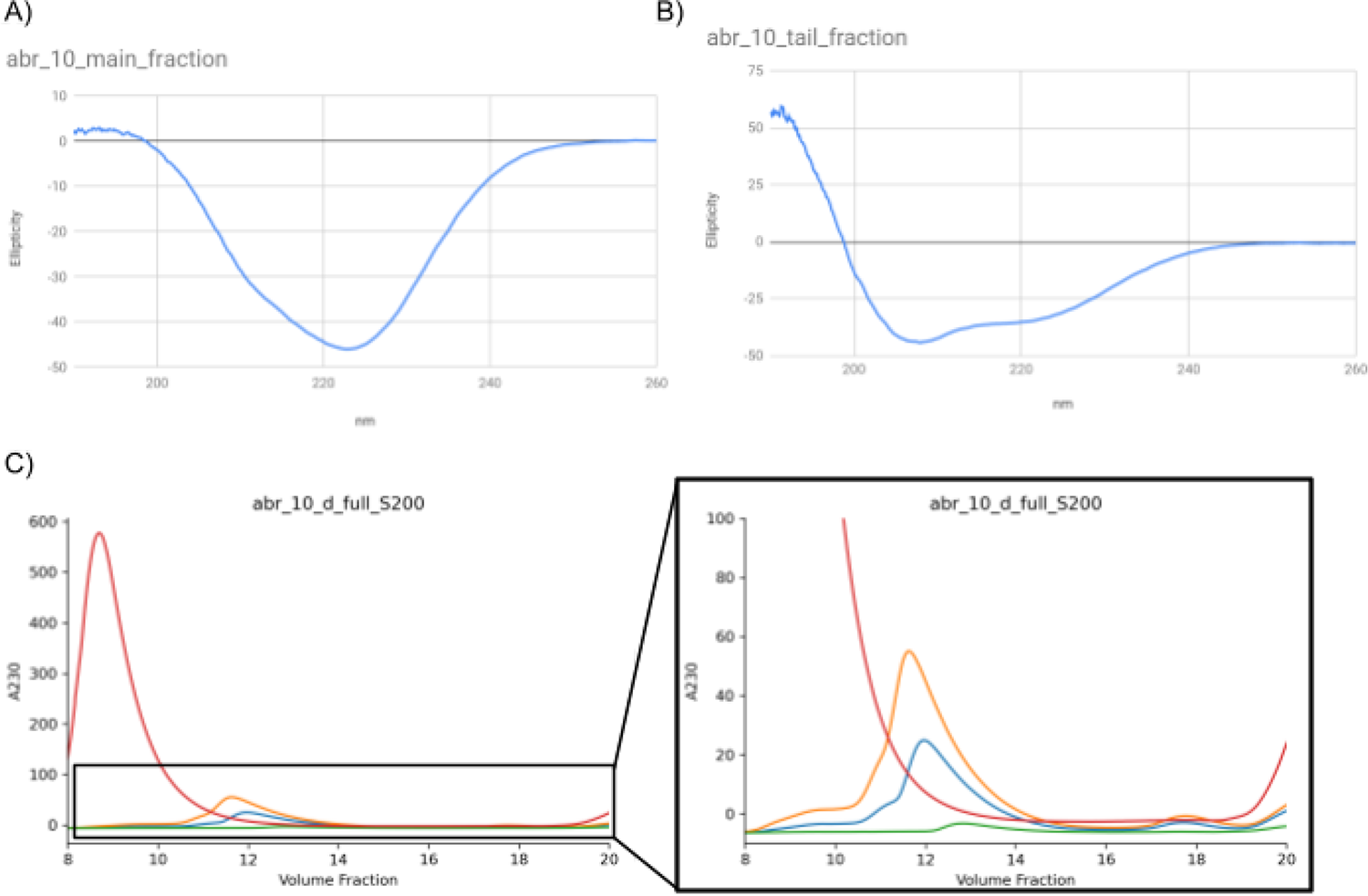
ABR 10 concentration dependent changes to SEC retention volume and CD trace profile. A) and B) show CD traces at ambient temperature for sample from the highly concentrated 8ml retention volume fraction and the dilute 12ml retention volume fraction. C) SEC profiles of a concentrated clarified lysate and serial dilutions (5X orange, 10X blue and 100X green dilutions relative to the original concentration in red). Inset shows what appears to be reversible soluble aggregation, with identifiable peaks at 9ml, 11ml, 12ml and 13 ml.

**Figure S.6.**
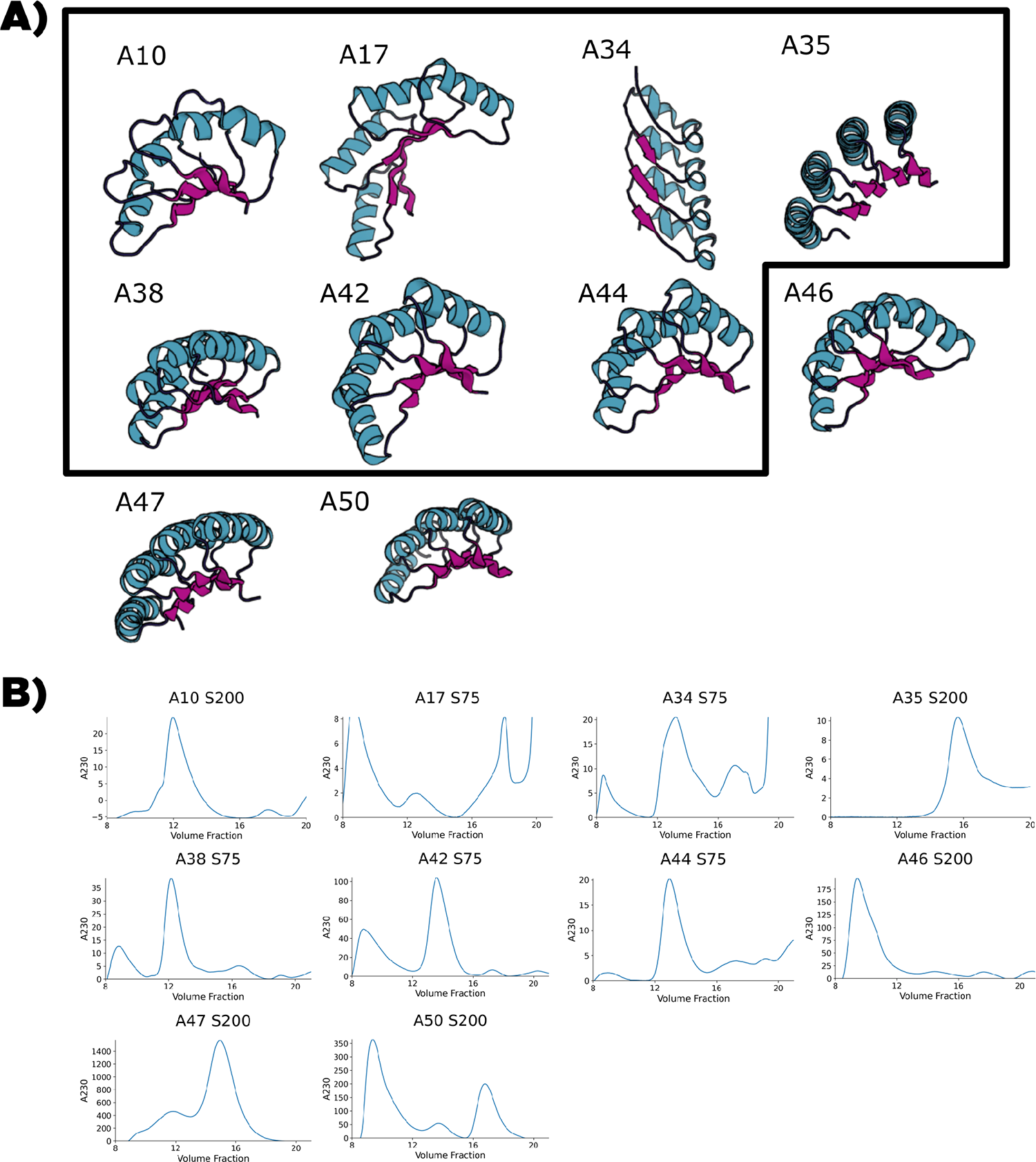
Rosetta-Based Uncapped designs Select Rosetta-based designs with tolerable solubility characteristics or interesting backbone features. A) backbone cartoon models with strands in pink and helices in teal. The black border outlines designs from design group 1 (minimal alpha-carbon constraints). The designs outside the border use both alpha-carbon and beta-carbon constraints. B) SEC traces from representative purification runs. For all models, a retention volume of approximately 14ml on S75 and one of 16ml on S200 corresponds roughly to a monomeric species. While these models had the best solubility characteristics in these design groups, they showed some oligomerization and concentration limited solubility.

